# pipemake: A pipeline creation tool using Snakemake for reproducible analysis of biological datasets

**DOI:** 10.1101/2024.12.20.629758

**Authors:** Andrew E. Webb, Scott W. Wolf, Ian M. Traniello, Sarah D. Kocher

**Affiliations:** Lewis-Sigler Institute for Integrative Genomics, Princeton University, Princeton, NJ, USA; Department of Ecology and Evolutionary Biology, Princeton University, Princeton, NJ, USA; Howard Hughes Medical Institute, Chevy Chase, MD, USA; Research Computing, Princeton University, Princeton, NJ, USA

## Abstract

The exponential growth in biological data generation has created an urgent need for efficient, reproducible computational analysis workflows. Here, we present pipemake, a computational platform designed to streamline the development and implementation of efficient and reproducible Snakemake workflows. pipemake creates modular pipelines that can be seamlessly integrated or removed from the platform without requiring reconfiguration of the core system, enabling flexible adaptation of workflows to different analytical needs across diverse fields. To demonstrate the platform’s capabilities, we created and implemented pipelines to reanalyze two distinct biological datasets. First, we recreated a population genomics analysis of the socially flexible halictid bee, *Lasioglossum albipes*, using pipemake-generated workflows for *de novo* genome annotation, processing of variant data, dimensionality reduction, and a genome-wide association study (GWAS). We then used pipemake to analyze behavioral tracking data from the common eastern bumble bee, *Bombus impatiens*. In both cases, pipemake workflows produced results consistent with published findings while substantially reducing hands-on analysis time. Overall, pipemake’s modular design allows researchers to easily modify existing pipelines or develop new ones without software development expertise. Beyond streamlining workflow creation, pipemake leverages the full Snakemake ecosystem to enable parallel processing, automated error recovery, and comprehensive analysis documentation. These features make pipemake an efficient and accessible solution for analyzing complex biological datasets. pipemake is freely available as a conda package or direct download at https://github.com/kocherlab/pipemake

## Introduction

The rate of big data production in biological science has rapidly increased over the past decades, leading to a discrepancy between the amount of data generated and our ability to meaningfully analyze it [1]. For example, advances in molecular biology have enabled high-throughput genomic data generation for nearly any organism, leading to a growing demand for genome-scale analyses in many different species. This has generated a significant need for approachable software and strategies to guarantee that findings are interpretable and reproducible [2, 3]. As such, there has been a massive influx of software packages designed by computational biologists targeted at addressing questions in the life sciences.

Much of the currently available software can be separated into three categories: toolkits, wrappers, and pipelines.

- **Toolkits** provide a collection of analytical tools that are typically designed to offer a variety of functions that operate on a specific file format and/or datatype (e.g. VCFtools, BCFtools, BEDTools, PLINK, GATK, SAMtools, etc.) [4, 5, 6, 7, 8]. While highly useful and invaluable in their area of expertise, they are unlikely to encompass the entirety of an analysis.
- **Wrappers** are software designed to call other libraries or executables (e.g. admixr, Funannotate, Galaxy, etc.) [9, 10, 11]. Often they involve simplifying the user interface and/or enabling the integration of software across multiple programming languages. Wrappers are typically used to perform a single task or command rather than a complete pipeline. While software wrappers are highly useful, they often lack features of the wrapped software and possess limitations imposed by method/language used to call the software.
- Self-contained **pipelines** have become more prevalent since the advent of conda environments and docker containers. They typically operate by automatically invoking commands from a collection of tools/software from different sources and/or programming languages (e.g. BRAKER, insectOR, etc.) [12, 13]. While such pipelines have the advantage of being easy to use, they may also act as a “black box” to most users who do not know the commands, their significance, and/or the order in which they operate. Additionally, the operations of contained pipelines are typically fixed and cannot be easily edited or repurposed.

Modern analyses typically require a combination of these different types of tools to operate. These tools are often combined by two key approaches. The first and far more common approach is to create lists of commands that call the necessary software. This method is inherently limiting due to hard-coded filenames and arguments that quickly make long lists of commands impractical to modify or operate with larger datasets and more sophisticated pipelines. These limitations frequently lead to the second approach, in which the software is repackaged within a new wrapper or pipeline. While the latter approach is often tempting, it may yield an overly complicated command structure and potentially limited, inefficient, and/or confusing resource requirements.

The limitations of both approaches can be overcome by using a workflow management system, such as Snakemake [14]. Snakemake creates workflows using short, text-based files that contain a standardized rule system. Each rule represents a single procedure within a workflow, and rules will be automatically invoked in a user-defined order to produce the desired output. Rules may vary in required resources such as threads and memory, file formats/types, and even software versions, all of which are customizable by the user. Rules may also be run in parallel, expediting workflows by fully utilizing High Performance Computing (HPC) environments or even systems with higher core counts. Snakemake also offers an ecosystem that is easily configurable, editable, and more importantly, reproducible.

While Snakemake provides an attractive ecosystem, some issues remain. First, developers are required to learn the rule nomenclature, which may present a challenge to those unfamiliar with programming. Additionally, reconfiguring and/or repurposing Snakemake rules can be unclear and error-prone for inexperienced developers. For researchers who are only interested in running a standalone analysis, such obstacles may result in Snakemake being an impractical solution.

For users more comfortable within the Snakemake ecosystem, some inconveniences still remain. Configuration files (“configs”) mitigate some of the issues in reconfiguring Snakemake workflows, but there is no system to rapidly generate or modify configs beyond editing previously created ones. While this is not inherently problematic, it represents a substantial investment of time, is highly error-prone, and is more likely to result in loss or corruption of the original config. Snakemake files may be reused across workflows, but this process is similarly subject to coding errors, especially if similar files with key parameter changes exist in a user’s file system. Moreover, workflow integrity is compromised without robust record-keeping and easy access to file modification history. This creates a need for a straightforward means of implementing a workflow manager, like Snakemake, within a platform that transparently manages the more difficult aspects of workflow generation, modification, and execution.

To this end, we present pipemake as a solution to reduce the barrier for users to generate and run reproducible computational analyses in the Snakemake ecosystem. pipemake is a pipeline creation tool designed to facilitate the development of Snakemake workflows to optimize computational efficiency and reproducibility. The workflows produced with pipemake can interface effectively with high performance computing systems to handle resource allocation using Snakemake’s built-in capabilities. Thus, pipemake enables the parallel processing of tasks and automatic determination of the optimal execution order. Moreover, workflows may be automatically resumed from a point of failure without a need to rerun previously completed steps.

pipemake was designed to streamline pipeline development by creating a flexible, modular platform with swappable pipelines that can easily reuse previously written Snakemake code. The package includes a collection of curated, customizable genomic and behavioral analysis pipelines for researchers seeking to rapidly integrate Snakemake-based workflows into their research, and additional modules can easily be integrated into the platform. Instead of a collection of immutable pipelines, pipemake is a unique platform that provides a flexible system for simplifying the creation and modification of Snakemake workflows. This nature allows pipemake to support a larger community of researchers than if it concentrated on a single aspect alone. Our hope is to grow this community as more snakemake workflows are developed, especially from different fields of study.

To demonstrate the broad utility of pipemake, we replicated computational analyses from previous work in two distinct fields of study: a population genomics study of social behavior in a socially flexible sweat bee, *Lasioglossum albipes*, and an automated behavioral tracking experiment in the common eastern bumble bee, *Bombus impatiens* [15, 16]. The pipelines we developed successfully replicated findings from both studies, and since they were developed within the pipemake platform, they may be easily reused or repurposed for independent datasets.

### pipemake

#### Design

pipemake was designed to operate using two primary filetypes that we outline in the following sections: snakemake files and a pipemake config file.

### Snakemake files

pipemake uses standard Snakemake files, which maintain their primary function of including the necessary rule(s) for a procedure or analysis [14]. The only requirement imposed by pipemake is that all paths begin with a pipemake-specific configurable directory to ensure that files are correctly assigned. pipemake is also capable of using Snakemake files to determine the necessary requirements for configuration (i.e. parameters, containers, resources, paths, etc.) and confirm all requirements were provided by the user. While pipemake operates on standard Snakemake files, ideally they should be modular in function and have configurable paths and parameters. The primary benefit of modular Snakemake files within the pipemake platform is that they can be easily reused, with the only requirement being the consistent specification of filenames and directories (Fig. 1).

**Fig. 1:**
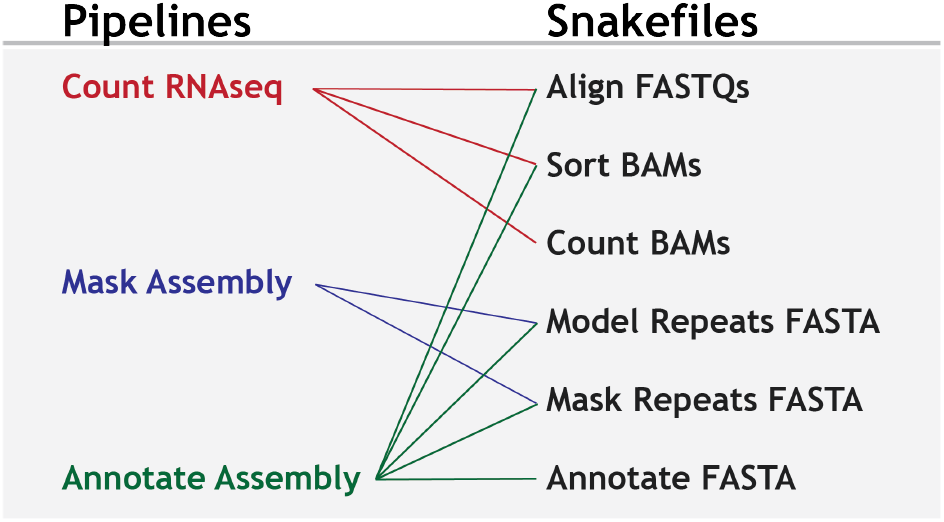
Example pipelines and Snakemake files. pipemake uses a simple system to associate Snakemake files with pipelines allowing them to be reused where appropriate.

Snakemake rules are also capable of operating within a conda environment or a singularity container [17, 18]. We chose to limit Snakemake rules in pipemake to singularity containers in order to reduce complexity and prioritize the benefits of providing all required software in container format. A key benefit of containers is that they enable software to operate within any operating system, a critical feature for maintaining the compatibility of the pipemake platform when adding future pipelines. Moreover, Snakemake automatically downloads and builds containers, eliminating the need for users to maintain software themselves [19].

To take full advantage of this functionality, we have created containers for all software currently required by our pipelines, and we will add more containers as new official pipelines are released. We have also designed pipemake to allow users to define a directory in which containers are stored. This option was added for groups who wish to have one set of containers that are shared by multiple users.

### Pipeline Configuration files

pipemake configuration files are YAML-based and are used by pipemake to define pipelines and provide: command-line arguments, the procedure to standardize input for the workflow, and a list of the required Snakemake files (Fig. 2). This is the primary file that users will need to define their own pipelines.

**Fig. 2:**
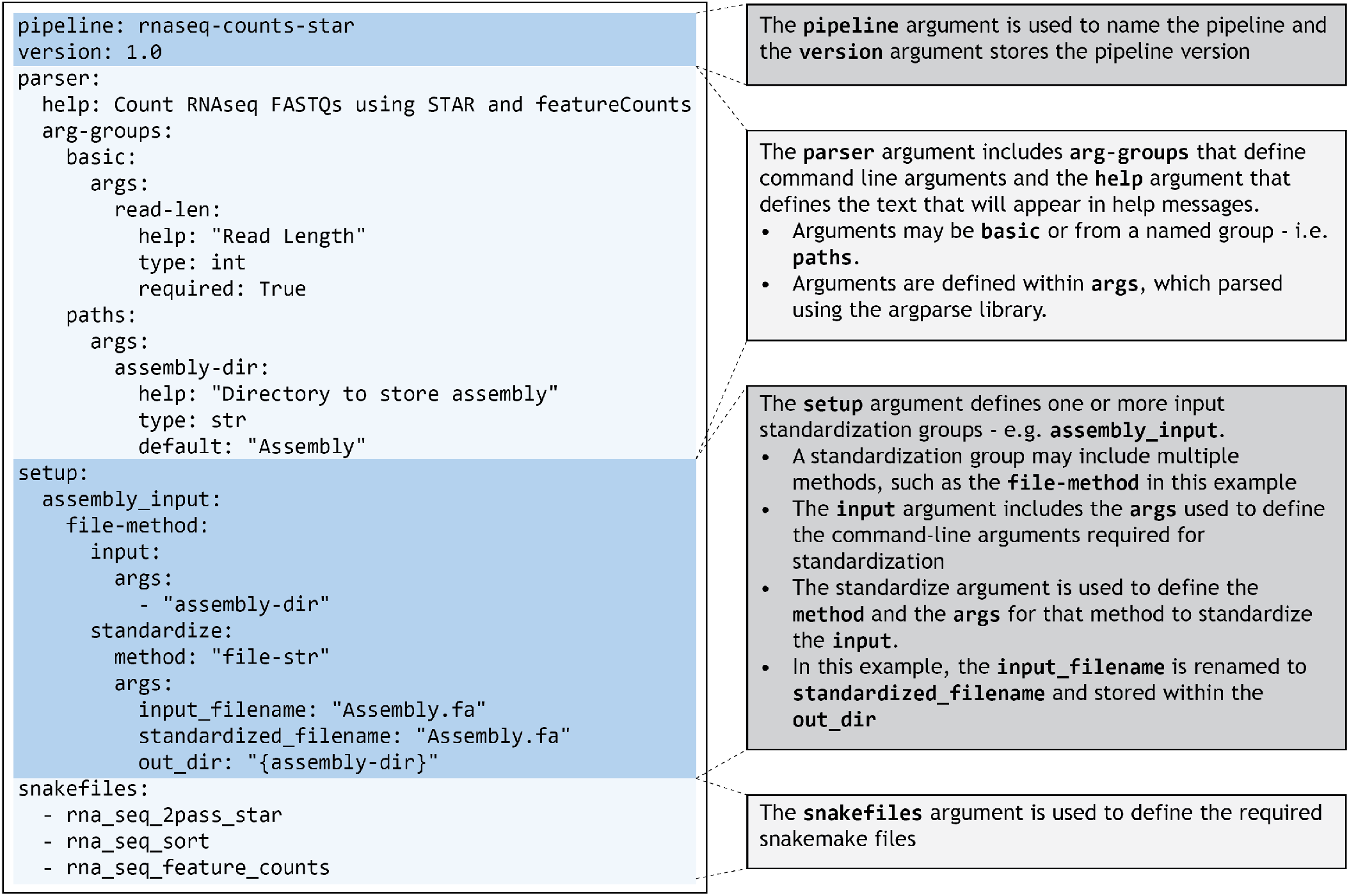
Structure of a Pipeline Configuration File. pipemake pipeline configuration files are separated into four categories: pipeline assignment, parser arguments, input standardization procedure, and Snakemake file requirements.

To define a pipeline, the only requirements are i) a unique pipeline name, ii) a description of the pipeline, and iii) the pipeline version. The complexity of the command-line arguments and input standardization is mostly dependent on the desired configurability of a pipeline. At a minimum, only the file input requires a command-line argument and a standardization procedure. Command-line arguments are parsed using the argparse python library and follow their standard nomenclature with some limitations to minimize compatibility issues. Standardization procedures are primarily used to ensure the input is correctly assigned for a procedure. Files are standardized by creating a copy or symbolic link of the file with a standardized name. It is also possible to have a standardization processes within a Snakemake file, such as providing a collection of sequence read archive (SRA) IDs which are standardized by directly downloading the files from the database [20]. And lastly, snakemake files are associated with a pipeline by including their filename and may be added or removed as needed.

A typical pipemake workflow begins by calling the executable, which will produce a list of the available pipelines along with a description of their function. Calling pipemake followed by the name of a pipeline will automatically produce the help text, listing all the relevant command-line arguments for the pipeline.

For most pipelines, the only command-line arguments required are the input (e.g. files, directories, SRA IDs, etc.) and critical parameters. When pipemake is called with a pipeline and the required command-line arguments, a workflow directory is produced along with the necessary workflow Snakemake files (Fig. 3).

**Fig. 3:**
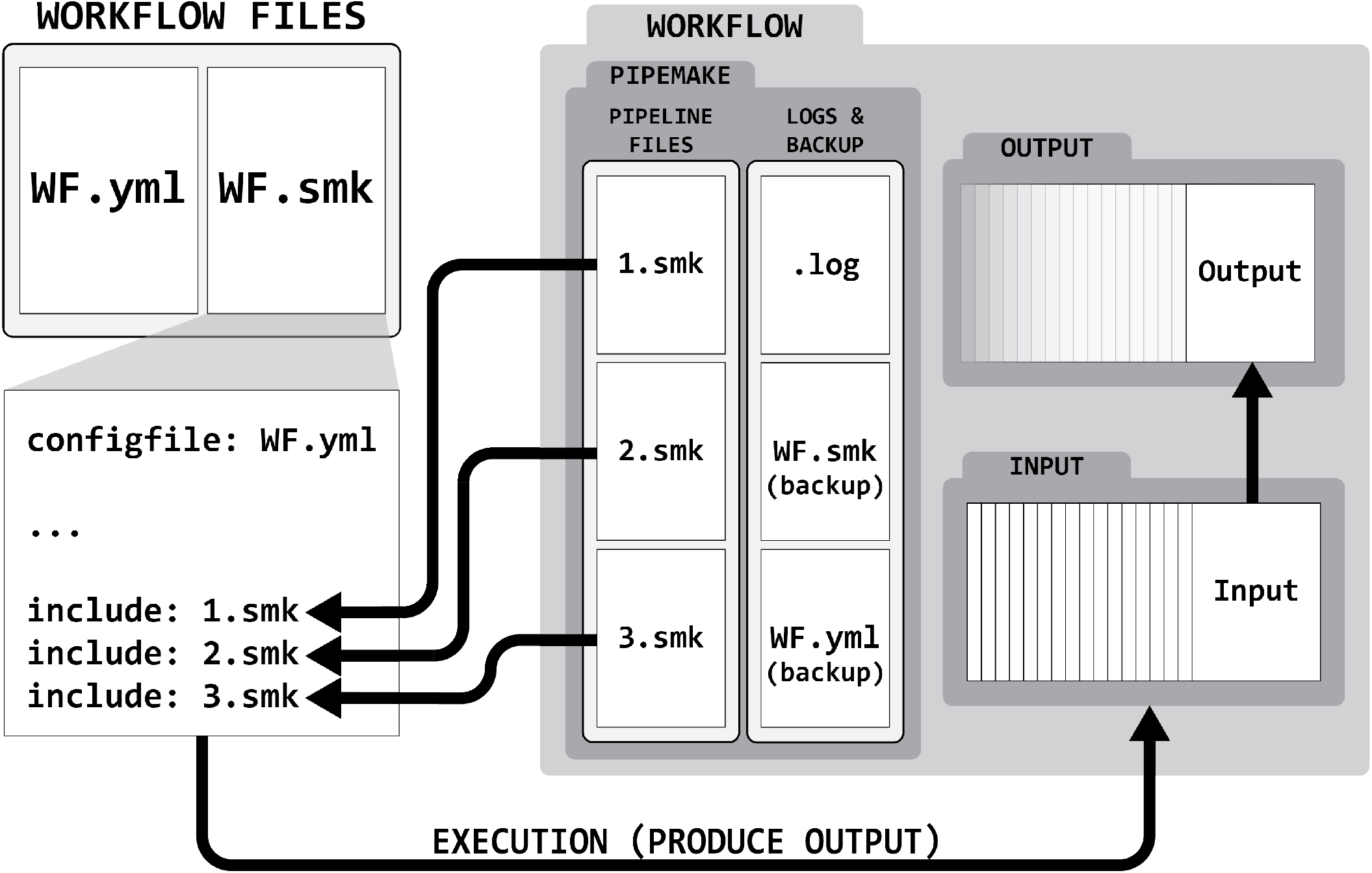
Structure of a pipemake Workflow. A pipemake workflow includes a directory, a Snakemake file, and a configuration file. The workflow Snakemake file is designed to be a single file that loads the config yml (WF.yml) and aggregates the rules defined within the pipeline Snakemake files (WF.smk). The pipeline Snakemake files (1.smk, 2.smk, 3.smk) are stored within the workflow directory and contain the rules for the workflow to function. When the workflow Snakemake file is executed it will produce the desired output within the workflow directory. The purpose of this schema is keep a record of all files required by the workflow to simplify record keeping and reproducibility. For this reason, pipemake automatically stores a backup of the workflow Snakemake and configuration files. Lastly, pipemake records all command-line arguments and file processing steps within a log file.

The workflow directory contains the pipemake directory and the standardized input or symbolic links, if not being downloaded from a database by the pipeline. The pipemake directory is used to store the pipeline Snakemake files, log files, and copies of the workflow Snakemake files. The workflow Snakemake file is used to define the desired output and links to the Snakemake files within the pipemake directory. A configuration file is also produced which defines all the required configurables alongside the default resources for each rule in the pipeline.

To start an analysis, the user is only required to execute Snakemake with the workflow Snakemake file. This allows users the flexibility of performing the analysis as desired - i.e. single-core, multi-core, multi-job on HPCs, etc. pipemake requires both the Snakemake files and pipeline configuration files to be stored in a predefined, configurable location. Additional pipelines may be added to pipemake simply by storing a new pipeline configuration file alongside any necessary Snakemake files. Once stored, pipemake will be capable of operating the new pipeline.

Several pipelines are included in the current release that enable a range of data analyses, including applications for: *de novo* genome annotations, the analysis of transcriptomic and chromatin accessibility data, population genomic data, and automated behavioral tracking data (see Table 1 for complete list). For detailed installation instructions and an example of pipemake implementation, please see the full documentation: https://pipemake.readthedocs.io/en/latest/.

**Table 1.** Current pipelines in pipemake.

| Pipeline | Function |
| --- | --- |
| annotate-braker3 | Annotate an assembly using BRAKER3 |
| annotate-braker3-with-bam | Annotate an assembly using BRAKER3 using a processed BAM file |
| annotate-genes-eggnog | Annotate genes using eggNOG |
| annotate-utrs-peaks2utr | Annotate an 3prime-UTRs using peaks2utr using a merged BAM file |
| atacseq-peaks-macs3 | Generate peaks for bulk ATAC-Seq data using bwa and MACS3 |
| codon-alignments | Create codon alignments from untranslated CDS multiple sequence files |
| fastq-filter | Filter FASTQ files using fastp |
| filter-model-vcf | Filter a model in resequencing VCF files using bcftools |
| filter-pops-vcf | Filter populations in resequencing VCF files using bcftools |
| filter-vcf | Filter resequencing VCF files using bcftools |
| mask-assembly | Annotate an assembly using BRAKER3 |
| reseq-calc-nsl | Calculate nSL using selscan |
| reseq-calc-xpnsl | Calculate XP-nSL using selscan |
| reseq-popgen | Run basic PopGen analyses on resequencing data |
| rnaseq-counts-star | Count RNAseq reads within a genome assembly using STAR and featureCounts |
| seperate-pop-vcfs | Create separate VCF files for each population in a resequencing VCF file |
| tracking-naps | Track MP4 videos using SLEAP and NAPS |
| unmapped-fastqs-star | Create unmapped FASTQs using STAR |

#### Case study 1: pipemake implementation of *de novo* genome annotation

pipemake’s short-read annotation pipeline, `annotate-braker3`, operates using BRAKER3, HISAT2, and RepeatModeler [13, 21, 22, 23]. We evaluated the pipeline using a recently released chromosomal-level assembly of *Lasioglossum albipes*, 40 paired-end RNAseq samples (NCBI SRA accession: PRJNA1142947), and a homology database constructed from OrthoDB and the OMA Orthology database [24, 25, 26]. Our snakemake workflow completed in approximately 38 hours running on a maximum 120 cores (or 20 jobs) without any intervention.

The pipeline resulted in 10,979 genes with a BUSCO score of 97.7%. This is fewer than the 15,905 genes inferred from a previous, more fragmented assembly [15]. Therefore, this discrepancy is likely partially due to less fragmented gene models, which might also explain the lower BUSCO score of 95.6% of the previous annotations.

We found the `annotate-braker3` pipeline to be simple to operate and efficient with minimal investment by the user. The pipeline was also simple to scale, allowing for the number of samples to have minimal effect on the overall runtime. These strengths make `annotate-braker3` an ideal method for exploratory studies and a foundation from which more sophisticated methods could be developed.

#### Case study 2: pipemake can analyze population genetic data

To evaluate the ability of pipemake’s population genomics pipelines to streamline an analysis, we sought to replicate results from a previously population genomic study comparing social and solitary populations of a socially flexible sweat bee, *Lasioglossum albipes* [15] (NCBI PRJNA413432). Since the original publication, an improved chromosomal-level assembly has been released [24]. We used the updated assembly described above and previously published whole-genome resequencing data from [15], which we mapped to the new assembly.

We used the snpArcher platform [27] to create the variant calls and generate the raw VCF file used in subsequent analyses. We next used the pipemake pipeline `filter-model-vcf` (Table 1) to filter the VCF file to 139 samples in our model using bcftools [7]. By default, `filter-model-vcf` will filter an input VCF/BCF in two steps: i) remove samples not specified within the model and ii) only include biallelic SNPs that pass the specified thresholds for MAF, quality, and missingness. pipemake also offers pipelines that will filter on all samples or on multiple populations/species. After filtration, we had a total of 4,159,171 SNPs; this is considerably higher than reported in Kocher et al. 2018 but not surprising considering we used a higher quality assembly with fewer gaps as well as a joint genotyping pipeline [15].

We next used the pipemake pipeline `reseq-popgen` (Table 1) to perform a basic set of analyses on the filtered VCF using bcftools, plink, and GEMMA [6, 7, 28]. This pipeline includes options for calculating Fst, LD-pruning the VCF, and performing a PCA and GWAS on the pruned dataset (Fig. 4A) (see Supplemental Methods for additional details). Our snakemake workflow completed in approximately 7 minutes running on a maximum 200 cores (or 60 jobs) without any intervention.

**Fig. 4:**
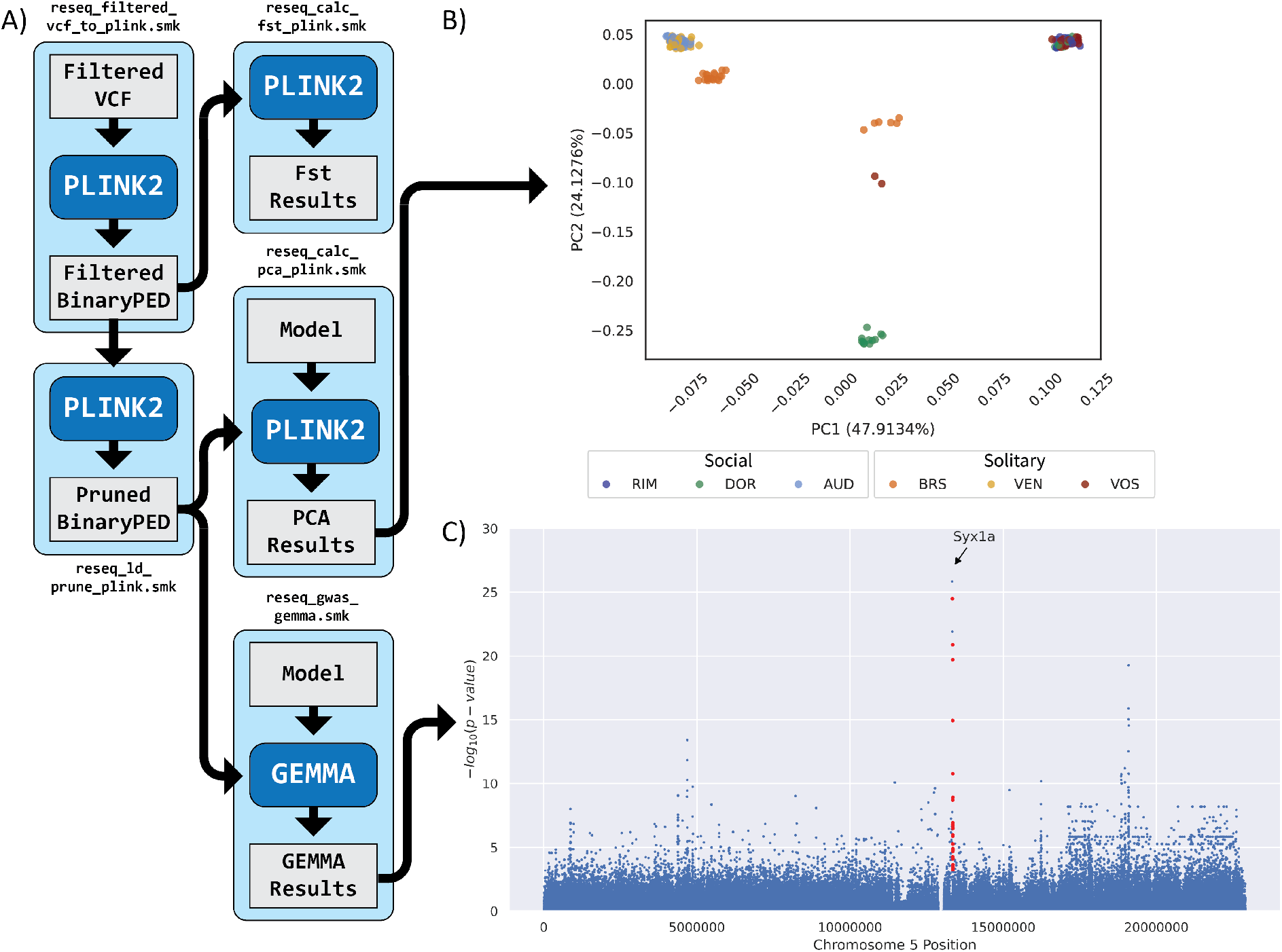
pipemake workflow for implementing Fst, principal component analysis (PCA), and a genome-wide association study (GWAS) using example data from the socially polymorphic sweat bee, *Lasioglossum albipes*. A) pipemake can implement genome-wide data analyses, such as the genome-wide PCA and GWAS using the `reseq-popgen` pipeline. The diagram depicts the primary snakemake files (light blue boxes) and the order in which they are executed to perform the analyses. Each snakemake file indicates the required input (light gray), the primary program (dark blue), and the output (light gray). Arrows indicate the flow from input to output, including output files that become input for subsequent snakemake files. Please note that not all steps are given for each snakemake rule for clarity. B) The PCA implemented by `reseq-popgen` replicates observations from [15]. For clarity, we have included a jitter to make each population visible among the clusters. C) The GWAS implemented by identified a similar peak associated with variation in social behavior in the sweat bee, *Lasioglossum albipes*. SNPs associated with social behavior and their genomic coordinates are located on chromosome 5 in an updated genome assembly. Each point in the manhattan plot represents a single SNP and its *−*log_10_ p-value. SNPs within 10kb of Syx1a and a FDR *<*0.2 are shown in red.

Overall, we found a large degree of overlap between Kocher et al 2018 and our reanalysis with pipemake. The PCA identified similar patterns of population clustering to Kocher et al. 2018, with the only exception being the separation of two samples from the VOS population from a cluster of samples from the BRS population (Fig. 4B). This is likely due to our use of a higher quality assembly and/or the improved genotyping calls from the updated joint genotyping pipeline. Moving on to the GWAS analysis, our pipeline replicated the peak at the location of *syntaxin 1A* (*syx1a*) (Fig. 4C). Closer examination of the peak found the seven variants, three within the first intron and four upstream, the same as reported in Kocher et al. 2018. In addition to the peak associated with *syx1a*, we also identified new peaks that were not observed in the original analysis and genome assembly (Fig. S1). Some of these peaks were found among odorant receptors, which suggests additional targets for future investigation. Overall, these findings illustrate how pipemake can successfully recapitulate previously published pipelines and results.

#### Case study 3: pipemake can perform automated behavioral tracking using NAPS

To demonstrate the versatility of the pipemake platform and its application beyond genomic datasets, we built and deployed a pipeline to replicate findings from a behavioral study of the common eastern bumble bee *Bombus impatiens*, recreating an analysis performed by NAPS (NAPS is ArUco Plus SLEAP), an automated behavioral monitoring software [16]. Specifically, NAPS utilizes hybrid tracking, or the coupling of a fiducial marker (i.e., a barcode) with pose estimation, a process in which a computer is tasked with recognizing an animal’s position in space [29, 30]. With NAPS, an investigator can monitor and quantify physical behaviors while also retaining individual identity over the course of an experiment.

The original NAPS workflow in Wolf et al. (2023) required video decoding, pose estimation, identity detection, and various post-processing steps to be run in sequence with frequent command-line intervention from the user. Using pipemake, these tasks were run in parallel and within a contained workflow that required minimal user intervention upon initiation, and our reanalysis perfectly matched initial findings: while either temporal tracking with SLEAP or hybrid tracking with NAPS identified nearly every individual in a video frame (Fig. 5, left), only NAPS correctly estimated the number of unique individuals filmed over the course of the tracked experiment (Fig. 5, right). Behavioral experiments involving different organisms or filming setups can reuse or adapt this workflow by swapping components like pose estimation models or behavioral analysis modules to best fit the focal system.

**Fig. 5:**
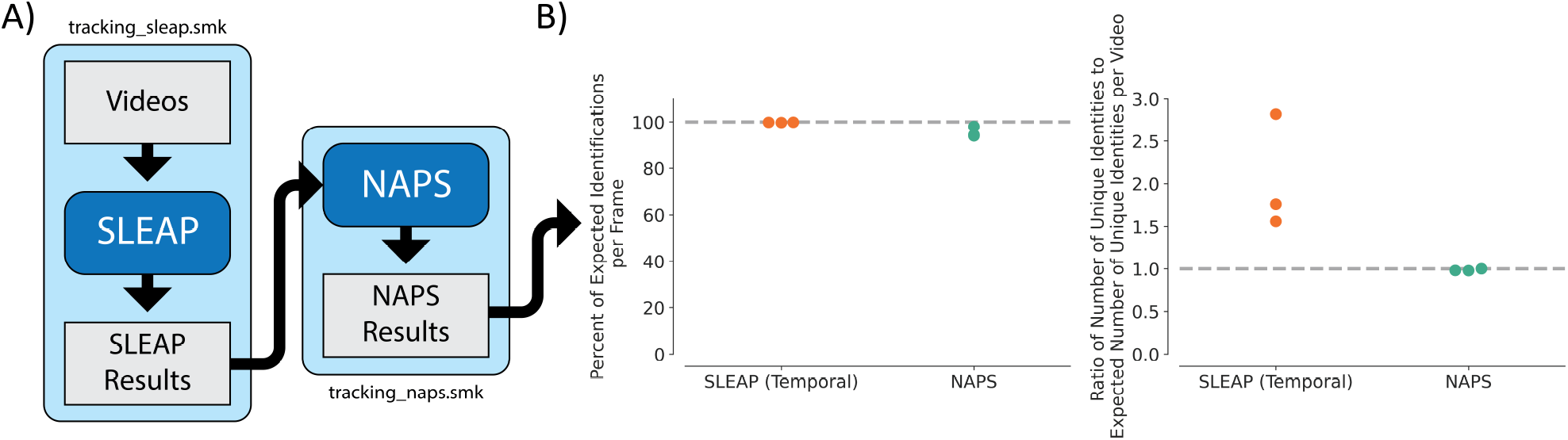
A comparison of the percent of expected identifications per frame between SLEAP an NAPS in an example data set. For our data set, we expect all 50 bees to be identified by SLEAP. For NAPS and ArUco, we expect that all active individuals, where the ArUco tag should be detected at some point throughout the video, to be detected. This number varies from 44 to 48 across three videos in the data set. (right) A comparison of the ratio of the expected number of unique identities compared to the expected number of unique identities per video. We expect 50 unique identities for SLEAP and from 44 to 48 for NAPS based on the number of active individuals in each video. These analyses were implemented using the `tracking-naps` pipeline in pipemake.

## Discussion

pipemake streamlines and expedites the development of transparent and reproducible Snakemake workflows for users with a wide range of programming experience. pipemake may be applied to any analysis pipeline requiring multi-step data processing workflows. pipemake’s modular structure makes it easy to extend or adapt workflows, allowing researchers to incorporate or modify existing pipelines without significant reconfiguration.

pipemake provides an accessible platform that can be used across a wide range of disciplines and input datasets. Here, we demonstrate cross-discipline generalizability by reanalyzing both genomic data as well as automated behavioral tracking data through a pipemake implementation of each dataset’s associated computational platforms. In both cases, pipemake replicated our previously published results, demonstrating its utility in building reproducible scientific data analysis.

The most obvious benefit of using pipemake is the ease of performing the same analysis on multiple species, across independent datasets, and in independent directories by only needing to change the relevant input arguments. For example, in comparative genomics, users often create a new directory that implements an identical file structure for each species of interest; with pipemake, a user may perform the same comparative analyses by simply modifying the name and location of the data input directory on the command-line.

Modifying existing pipelines and developing new ones is greatly simplified given pipemake’s modular nature. New pipelines can be readily created using a combination of existing Snakemake modules and new ones. For instance, the creation of our genome annotation pipeline was greatly streamlined as we integrated previously developed Snakemake modules for RNAseq alignment and repeat masking. Additionally, the creation of new pipeline configuration files is simplified by inserting the relevant components from previously created pipelines that use the same Snakemake files.

An unexpected benefit of pipemake’s design was the the ease of adding quality control (QC) modules to existing pipelines. From our experience, once a QC module was designed for a particular datatype or procedure it could easily be propagated to all relevant pipelines. Even rule-specific or unique QC modules could be readily added to a pipeline without the need to alter other Snakemake files.

Implementing pipelines and troubleshooting is also streamlined within the pipemake platform. Using modular Snakemake files often allows errors to be easily attributed to a single rule, this is especially true when developing from an existing pipeline. Errors due to insufficient resource allocations can be resolved using pipemake’s built-in resource-scalars for widespread issues or modifying the workflow configuration file for rule-specific issues. These same tools were also useful to adjust resources to better utilize parallel processing and the larger RAM allocations allowed on HPC clusters. Also, if desired, the workflow directory allows for pipeline modifications to be explored without impact to the original pipeline.

The workflow directory is an ideal target for data preservation, as it stores all relevant pipeline files, the input, and the output. Preserving the workflow directory also provides other critical benefits. The directory may be used to produce a detailed report on all commands performed within a pipeline alongside a visual representation of the pipeline. This greatly simplified our record-keeping and allows for the records to be easily recreated if lost. The directory may also be used to easily rerun a pipeline, greatly simplifying the process of replicating an analysis.

The primary goal our pipeline management system is to simplify the process of maintaining and obtaining pipelines. Future updates to pipemake will primarily focus on feature improvements that facilitate pipeline development and availability. We will begin by developing a graphical user interface (GUI) to simplify pipeline creation and reduce the barriers to creating pipeline configuration files. We are also exploring the practicality of implementing an online database to store and maintain pipelines.

pipemake’s modular nature and its ability to handle complex workflows make it an ideal platform to rapidly develop Snakemake workflows. Workflows designed in pipemake are scalable, easily reproducible, and streamlined for record-keeping and data preservation. Here, we have demonstrated pipemake’s potential by replicating analyses in two distinct disciplines of biology, however, the platform can be expanded to any application that can operate within a workflow. pipemake is freely available as a conda package or direct download at https://github.com/kocherlab/pipemake.

## Author contributions statement

AEW conceived of and wrote pipemake with input from SDK. SWW and IMT contributed pipemake modules and provided feedback on the software. AEW and SWW conducted data analyses. AEW wrote the original manuscript draft and received feedback from all co-authors.

## Acknowledgments

The authors thank Brian Arnold, Sarthok Rahman, and other members of the Kocher Lab for beta testing and input. This work is supported in part by funds from the National Institutes of Health (NIH DP2 GM137424), and by the Packard Foundation and Pew Biomedical Scholars Program. IMT is a Lewis-Sigler Scholar and SDK is an HHMI Freeman Hrabowski Scholar.

