## Supplementary Methods for "pipemake: A pipeline creation tool using Snakemake for reproducible analysis of biological datasets"

### Annotation of *Lasioglossum albipes*

The pipemake pipeline `annotate_braker3` requires: i) a FASTA-formatted assembly, ii) RNAseq FASTQs from one or more samples (both paired-end and single-end are allowed), and a FASTA-formatted file consisting of protein sequences from related species. The pipeline begins by aligning the FASTQs to the assembly using HISAT2, then masking the assembly using both RepeatModeler and RepeatMasker Kim et al. [2019], Smit, Arian et al. [2013], Flynn et al. [2020]. These processes may be done in parallel but both must be finished prior to annotating the assembly using BRAKER3 Gabriel et al. [2024]. To annotate *Lasioglossum albipes* we selected an unpublished chromosomal-level assembly, 40 paired-end RNAseq samples, and a protein sequence database constructed from the amino acid sequences of *Apis mellifera* (AMEL_HAv3.1), *Bombus impatiens* (BIMP_2.2), ortholog groups with an arthropod ortholog from OMA, and the arthropods database from odb11 Wallberg et al. [2019], Sadd et al. [2015], Altenhoff et al. [2024], Zdobnov et al. [2021].

In the following command we built the snakemake workflow by calling the assembly fasta `LALB_genome_v3.fasta.gz`, a RNAseq sample file `LALB_genome_v3.fasta.gz`, and the protein sequence database `ProteinHints_proteins.11.fasta`. We also used the optional arguments that define the species name, assembly and annotation version, and the workflow naming scheme.

```
pipemake annotate-braker3 --rnaseq-table LALB_Filtered_FASTQs.tsv --assembly-fasta
    LALB_genome_v3.fasta.gz --hints-fasta ProteinHints_proteins.11.fasta --busco-
    database hymenoptera_odb10 --species LALB --assembly-version v3 --annotation-
    version 1 --work-dir LALB_v3.1 --snakemake-job-prefix LALB_v3.1
```

Once the workflow was created we ran it using:

```
snakemake -s LALB_v3.1.smk --use-singularity
```

### VCF construction

The joint-genotyping pipeline of snpArcher requires: i) a FASTA-formatted assembly and ii) resequencing FASTQs from one or more samples which may be provided locally or via SRA IDs Mirchandani et al. [2024], Wheeler et al. [2007]. We selected a collection of *Lasioglossum albipes* samples from three social populations (AUD, DOR, RIM) and three solitary populations (VOS, BRS, VEN) originally collected for Kocher et al. 2018 Kocher et al. [2018]. We genotyped the samples using the *Lasioglossum albipes* assembly described in our annotation pipeline. We ran snpArcher with the standard parameters provided in the snpArcher configuration file with the exception of disabling the missingness filter. We disabled missingness due to the filter being implemented in pipemake with greater functionality.

snpArcher was ran using the following command:

```
snakemake -s LALB_v3.smk --use-conda
```

### VCF filtering

The pipemake pipeline `filter-model-vcf` requires: i) a VCF, ii) a model file, and iii) a model name. The model file is JSON-based file originally described in the Popgen Pipeline Platform

Webb et al. [2021]. A model file consists of one or more models, a model is used to store individual/sample relationships. For instance, one model may store the population-level relationship for each sample (i.e. sample A is from AUD) whereas another may store a categorical relationship such as sociality instead (i.e. sample A is social). `filter-model-vcf` filtered the output of snpArcher, `LALB_v3_raw.vcf.gz`, using a two-step procedure. It began by removing samples not found among the 139 sample specified within our sociality model, this was primarily used to remove samples with sequencing issues and inconsistent records. It next filtered the VCF to only include biallelic SNPs that passed the following thresholds: MAF $\geq$ 0.05, missingness $\leq$ 0.11, and QUAL $\geq$ 30. These thresholds were selected to match (or be equivalent to) those used in Kocher et al. 2018 Kocher et al. [2018]. The `filter-model-vcf` workflow was created using the following command:

```
pipemake filter-vcf --reseq-vcf LALB_v3_raw.vcf.gz --model-file LALB.model --model-name
     LALB --missing-cutoff 0.11 --species LALB --assembly-version v3 --work-dir
    LALB_v3_Filter --snakemake-job-prefix LALB_v3_Filter
```

Once the workflow was created we ran it using:

```
snakemake -s LALB_v3_Filter.smk --use-singularity
```

### PCA and GWAS analyses

The pipemake pipeline `reseq-popgen` requires: i) a VCF, ii) a model file, and iii) one or more model names to calculate Fst using PLINK2, perform a PCA using PLINK2, and perform a GWAS using GEMMA Purcell, Shaun and Chang, Christopher, Zhou and Stephens [2012]. The pipeline automatically creates a pruned dataset for the PCA and GWAS analyses using PLINK2. In general, each analysis will remove samples not found among the model and account for sample populations/categories. For instance, the sociality model includes two categories, social and solitary, when calculating Fst a single pairwise comparison is reported. In contrast, when calculating Fst using the population model, which consists of AUD, DOR, RIM, VOS, BRS, and VEN, 15 pairwise comparison are reported. This also holds among visualization where appropriate, for instance, the PCA analysis will automatically produce a plot of PC1 and PC2 with samples assigned to their respective category. We used the default settings for both the GWAS and PCA analyses, we pruned the VCF using a window size of 50 SNPs, a step size of 5, and r2 threshold of 0.8. The `reseq-popgen` workflow to created using the filtered VCF `LALB_v3.filtered.vcf.gz` and both the sociality and population models using the following command:

```
pipemake reseq-popgen --reseq-vcf LALB_v3.filtered.vcf.gz --model-file LALB.model --
    models LALB_Sociality LALB_Pops --ld-threshold 0.8 --species LALB --assembly-
    version v3 --work-dir LALB_v3_PopGen --snakemake-job-prefix LALB_v3_PopGen
```

Once the workflow was created we ran it using:

```
snakemake -s LALB_v3_PopGen.smk --use-singularity
```

Ben M Sadd, Seth M Barribeau, Guy Bloch, Dirk C De Graaf, Peter Dearden, Christine G Elsik, Jürgen Gadau, Cornelis Jp Grimmelikhuijzen, Martin Hasselmann, Jeffrey D Lozier, Hugh M Robertson, Guy Smagghe, Eckart Stolle, Matthias Van Vaerenbergh, Robert M Waterhouse, Erich Bornberg-Bauer, Steffen Klasberg, Anna K Bennett, Francisco Câmara, Roderic Guigó, Katharina Hoff, Marco Mariotti, Monica Munoz-Torres, Terence Murphy, Didac Santesmasses, Gro V Amdam, Matthew Beckers, Martin Beye, Matthias Biewer, Márcia Mg Bitondi, Mark L Blaxter, Andrew Fg Bourke, Mark Jf Brown, Severine D Buechel, Rossanah Cameron, Kaat Cappelle, James C Carolan, Olivier Christiaens, Kate L Ciborowski, David F Clarke, Thomas J Colgan, David H Collins, Andrew G Cridge, Tamas Dalmay, Stephanie Dreier, Louis Du Plessis, Elizabeth Duncan, Silvio Erler, Jay Evans, Tiago Falcon, Kevin Flores, Flávia Cp Freitas, Taro Fuchikawa, Tanja Gempe, Klaus Hartfelder, Frank Hauser, Sophie Helbing, Fernanda C Humann, Frano Irvine, Lars S Jermiin, Claire E Johnson, Reed M Johnson, Andrew K Jones, Tatsuhiko Kadowaki, Jonathan H Kidner, Vasco Koch, Arian Köhler, F Bernhard Kraus, H Michael G Lattorff, Megan Leask, Gabrielle A Lockett, Eamonn B Mallon, David S Marco Antonio, Monika Marxer, Ivan Meeus, Robin Fa Moritz, Ajay Nair, Kathrin Näpflin, Inga Nissen, Jinzhi Niu, Francis Mf Nunes, John G Oakeshott, Amy Osborne, Marianne Otte, Daniel G Pinheiro, Nina Rossié, Olav Rueppell, Carolina G Santos, Regula Schmid-Hempel, Björn D Schmitt, Christina Schulte, Zilá Lp Simões, Michelle Pm Soares, Luc Swevers, Eva C Winnebeck, Florian Wolschin, Na Yu, Evgeny M Zdobnov, Peshtewani K Aqrawi, Kerstin P Blankenburg, Marcus Coyle, Liezl Francisco, Alvaro G Hernandez, Michael Holder, Matthew E Hudson, LaRonda Jackson, Joy Jayaseelan, Vandita Joshi, Christie Kovar, Sandra L Lee, Robert Mata, Tittu Mathew, Irene F Newsham, Robin Ngo, Geoffrey Okwuonu, Christopher Pham, Ling-Ling Pu, Nehad Saada, Jireh Santibanez, DeNard Simmons, Rebecca Thornton, Aarti Venkat, Kimberly Ko Walden, Yuan-Qing Wu, Griet Debyser, Bart Devreese, Claire Asher, Julie Blommaert, Ariel D Chipman, Lars Chittka, Bertrand Fouks, Jisheng Liu, Meaghan P O'Neill, Seirian Sumner, Daniela Puiu, Jiaxin Qu, Steven L Salzberg, Steven E Scherer, Donna M Muzny, Stephen Richards, Gene E Robinson, Richard A Gibbs, Paul Schmid-Hempel, and Kim C Worley. The genomes of two key bumblebee species with primitive eusocial organization. *Genome*

*Biology*, 16(1):76, December 2015. ISSN 1474-760X. doi: 10.1186/s13059-015-0623-3. URL `https://genomebiology.biomedcentral.com/articles/10.1186/s13059-015-0623-3`.

Adrian M Altenhoff, Alex Warwick Vesztrocy, Charles Bernard, Clement-Marie Train, Alina Nicheperovich, Silvia Prieto Baños, Irene Julca, David Moi, Yannis Nevers, Sina Majidian, Christophe Dessimoz, and Natasha M Glover. OMA orthology in 2024: improved prokaryote coverage, ancestral and extant GO enrichment, a revamped synteny viewer and more in the OMA Ecosystem. *Nucleic Acids Research*, 52(D1):D513–D521, January 2024. ISSN 0305-1048, 1362-4962. doi: 10.1093/nar/gkad1020. URL `https://academic.oup.com/nar/article/52/D1/D513/7420097`.

Evgeny M Zdobnov, Dmitry Kuznetsov, Fredrik Tegenfeldt, Mosè Manni, Matthew Berkeley, and Evgenia V Kriventseva. OrthoDB in 2020: evolutionary and functional annotations of orthologs. *Nucleic Acids Research*, 49(D1):D389–D393, January 2021. ISSN 0305-1048, 1362-4962. doi: 10.1093/nar/gkaa1009. URL `https://academic.oup.com/nar/article/49/D1/D389/5983625`.

Cade D Mirchandani, Allison J Shultz, Gregg W C Thomas, Sara J Smith, Mara Baylis, Brian Arnold, Russ Corbett-Detig, Erik Enbody, and Timothy B Sackton. A Fast, Reproducible, High-throughput Variant Calling Workflow for Population Genomics. *Molecular Biology and Evolution*, 41(1):msad270, January 2024. ISSN 1537-1719. doi: 10.1093/molbev/msad270. URL `https://doi.org/10.1093/molbev/msad270`.

D. L. Wheeler, T. Barrett, D. A. Benson, S. H. Bryant, K. Canese, V. Chetvernin, D. M. Church, M. DiCuccio, R. Edgar, S. Federhen, M. Feolo, L. Y. Geer, W. Helmberg, Y. Kapustin, O. Khovayko, D. Landsman, D. J. Lipman, T. L. Madden, D. R. Maglott, V. Miller, J. Ostell, K. D. Pruitt, G. D. Schuler, M. Shumway, E. Sequeira, S. T. Sherry, K. Sirotkin, A. Souvorov, G. Starchenko, R. L. Tatusov, T. A. Tatusova, L. Wagner, and E. Yaschenko. Database resources of the National Center for Biotechnology Information. *Nucleic Acids Research*, 36(Database):D13–D21, December 2007. ISSN 0305-1048, 1362-4962. doi: 10.1093/nar/gkm1000. URL `https://academic.oup.com/nar/article-lookup/doi/10.1093/nar/gkm1000`.

Sarah D. Kocher, Ricardo Mallarino, Benjamin E. R. Rubin, Douglas W. Yu, Hopi E. Hoekstra, and Naomi E. Pierce. The genetic basis of a social polymorphism in halictid bees. *Nature Communications*, 9(1):4338, October 2018. ISSN 2041-1723. doi: 10.1038/s41467-018-06824-8. URL `https://www.nature.com/articles/s41467-018-06824-8`.

Andrew Webb, Jared Knoblauch, Nitesh Sabankar, Apeksha Sukesh Kallur, Jody Hey, and Arun Sethuraman. The Pop-Gen Pipeline Platform: A Software Platform for Population Genomic Analyses. *Molecular Biology and Evolution*, 38(8):3478–3485, July 2021. ISSN 1537-1719. doi: 10.1093/molbev/msab113. URL `https://academic.oup.com/mbe/article/38/8/3478/6265476`.

Purcell, Shaun and Chang, Christopher. PLINK 2.0. URL `www.cog-genomics.org/plink/2.0/`.

Xiang Zhou and Matthew Stephens. Genome-wide efficient mixed-model analysis for association studies. *Nature Genetics*, 44(7):821–824, July 2012. ISSN 1061-4036, 1546-1718. doi: 10.1038/ng.2310. URL `https://www.nature.com/articles/ng.2310`.
