## Supplementary figures and images for "pipemake: A pipeline creation tool using Snakemake for reproducible analysis of biological datasets"

### Supplementary Figures

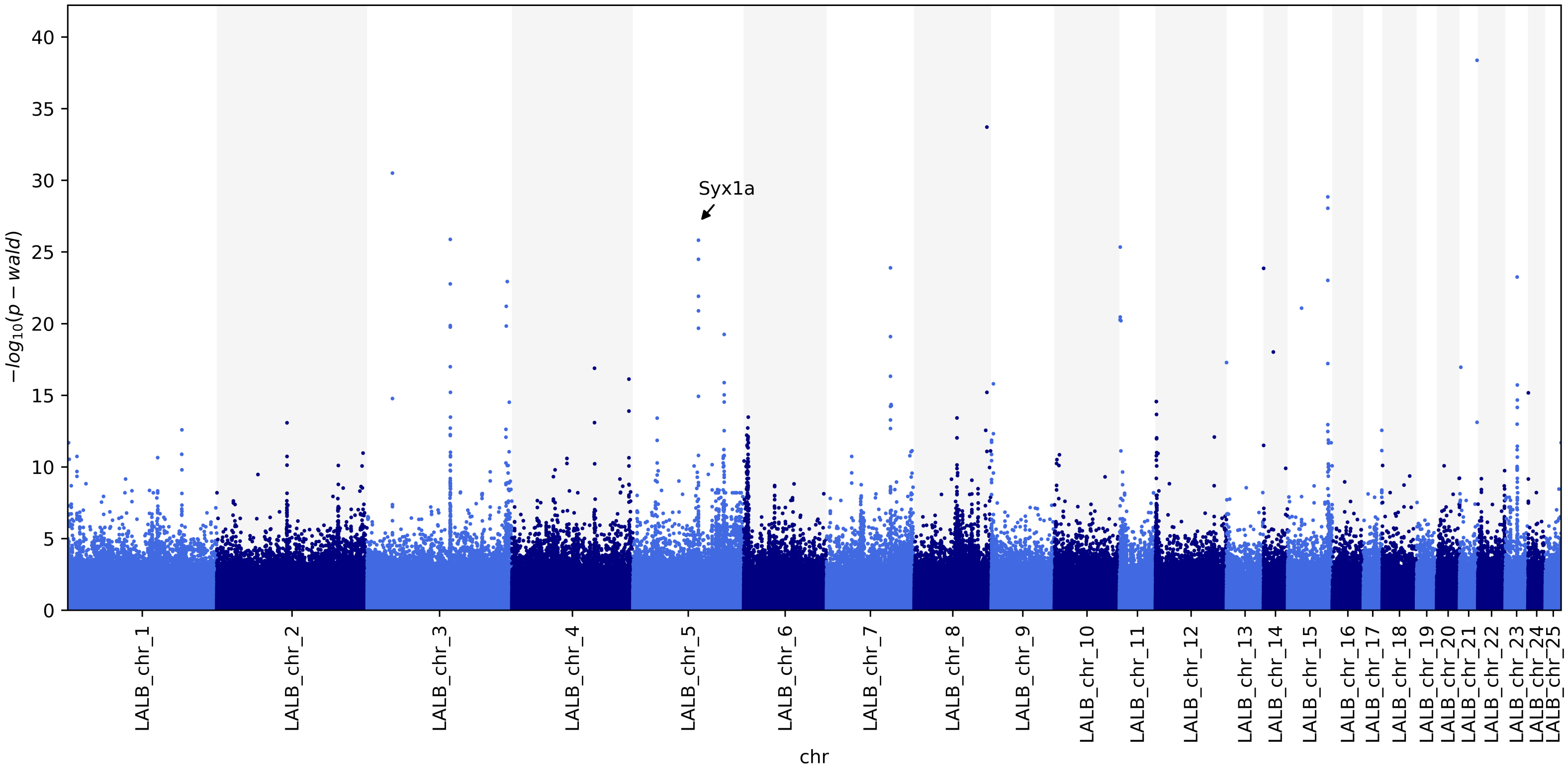


**Figure S1: GWAS replicates previous identification of Syx1a**
